# The Effects of Pressure and Temperature on the Thermodynamics of α-Helices

**DOI:** 10.1101/2025.01.29.635541

**Authors:** George I. Makhatadze, Caitlyn Moustouka, Carleton Coffin, Franco O. Tzul

**Author notes:** **Corresponding Author:** George I. Makhatadze.

## Abstract

In light of the recent realization that a large fraction of microbial biomass lives under high hydrostatic pressure, there is a renewed interest in understanding molecular details by which proteins in these organisms modulate their functional native state. The effects of pressure on protein stability are defined by the volume changes between native and denatured states. The conformational ensemble of the denatured state can depend on several extrinsic variables, such as pH and ionic strength of solvent, temperature, and pressure. The effect of the latter on the elements of the secondary structures, and α-helical structures has been inconclusive. This has been largely due to the inherent difficulties of high-pressure experiments. Here, we adapted the method of choice, circular dichroism spectroscopy, on a well-established series of model peptides to study helical structure formation, while focusing on the pressure and temperature dependencies of the helix-coil transition. We find that at low temperatures, pressure stabilizes the helical structure, suggesting that the volume of the helix-coil transition is positive. However, at higher temperatures (>40°C), the volume changes become negative, and pressure destabilizes the helical structure.

## Introduction

The thermodynamic stability of proteins is defined by the Gibbs free energy differences (Δ*G*) between the native, folded states and the denatured states. While the native state can be rather well characterized by various biophysical methods, the conformational ensemble of the denatured state and, thus, associated thermodynamics, remains elusive. In particular, the question of the residual structure in the denatured state ensemble has always been the subject of intense debates (see, e.g., ^1-6^). It was argued that the denatured state ensemble in high concentrations of urea or guanidine closely resembles the random coil state, while heat-denatured and, even more so the cold-denatured states, contain a significant amount of residual helical secondary structure ^7^. More recently, the effects of hydrostatic pressure have come to the attention of discussions of the residual structure in the denatured state ^8^. While it has always been accepted that pressure, together with temperature, are major intrinsic environmental variables, only recently has it been realized that a large fraction of life on Earth lives under excess pressure in the ocean and in the subsurface crevices ^9^. Thus, the stability of proteins from these organisms might be profoundly affected by the pressure. The effects of pressure on protein stability are defined by the sign of the volume change (Δ*V*) associated with unfolding: if Δ*V* is negative, an increase in pressure will lead to destabilization of the native state, whereas if Δ*V* is positive, an increase in pressure will lead to an increase in stability of the native state ^10^. The Δ*V* values for the elements of protein secondary structure, such as the β-hairpin and α-helix, have been measured experimentally and estimated computationally. Our direct experimental data on a Trp-zip peptide, forming a β-hairpin structure, shows positive Δ*V*, which was well predicted by our computational model ^10^. There are predictions of positive Δ*V* for the helix-coil transition based on several computational papers ^11-14^. Unfortunately, the experimental validation of helix stabilization has been based on indirect biophysical methods ^15-16^. The Δ*V* values measured for AK16 peptide are based on FT-IR spectroscopy ^16^, which requires spectral decomposition using model-dependent curve fitting to obtain the contribution of α-helix. The Δ*V* values for the AAQAA_3_ peptide were obtained using temperature jump (T-jump) and pressure jump (P-jump) monitored by triplet-triplet energy transfer in labeled peptide ^15^. Our own attempts to measure Δ*V* using pressure perturbation calorimetry (PPC) were not successful due to the low enthalpy of unfolding, resulting in a very broad transition not detectable by the changes in expansivity (Δ*α*) measured by PPC ^17^.

This notion of secondary structure stabilization under high-hydrostatic pressure would suggest that pressure-induced folding might lead denatured state ensembles to take on higher degrees of secondary structural content. We thus decided to directly characterize the helical content of peptides at high hydrostatic pressure and, by doing so, provide detailed experimental validation for the sign and magnitude of Δ*V* upon the helix-coil transition. To this end, we have used a high-pressure cell adapted to be used with one of the most well-characterized methods for experimentally assessing secondary structure in peptides and proteins: circular dichroism (CD) spectroscopy. These experiments also require a model host-guest peptide system that is relatively simple and offers a wide range of helical content under ambient pressure. Such a model system studied by us previously is the XEARA6 peptides, which are 32 amino acid-long peptides containing six repeats of the pentameric sequence XEARA (plus Y at the C-terminus to allow absorbance measurements for concentration), where the guest position X=G, A, V, L, I, or M. We have previously demonstrated that these synthetic peptides have varying degrees of helical structure at ambient pressure ^18-19^. The presence of the straight side-chain amino acid Arg, and γ-branched Glu in the Ala-rich sequence context favors helix formation ^20-24^. The I to I + 3 spacing of arginine and glutamate promotes stabilizing sidechain-to-sidechain interactions ^25^. The XEARA6 peptides show a significant variation in the fraction of helical structure as measured by far-UV CD spectroscopy ^18^. As expected, the GEARA6 peptide does not contain any helical structure as judged by the far-UV CD spectrum (Figure 4). The amount of helical structure in the rest of the peptides follows the trend expected based on well-established helical propensities of amino acid residues ^20-22, 26^. Peptides containing β-branched side chains in the X position (e.g., VEARA6 and IEARA6) have lower helical content than those with -branched (e.g., LEARA6) or unbranched (e.g., AEARA6) side chains.

To further validate the conclusions from high-pressure CD spectroscopy experiments, we also developed protein constructs that use the two peptides with the most extreme differences in helical propensity, GEARA6 and AEARA6, as linkers between two FRET-compatible fluorescent proteins to perform high-pressure fluorescence and small-angle x-ray scattering (SAXS) experiments. The results from these two orthogonal approaches affirm the results obtained from the CD experiments. The overall conclusion is that the volume changes upon helix-coil transition are positive at low temperatures. As temperature increases, the volume changes become smaller and even negative at temperatures above 40°C, leading to a destabilization of helical structure with an increase in pressure.

## Results and Discussion

The goal of this work was to probe the effect of pressure on the stability of α-helices. As a model system, we chose to use a series of XEARA6 peptides that have been previously well-characterized and shown to be an excellent model to study the thermodynamic stability of α-helices ^18-19^. This peptide model consists of six repeats (plus C-terminal Y for concentration measurements), with each repeat containing a guest position X. This ensures a robust effect of the substitutions on the structural properties of peptides. The XEARA6 peptides have a high fraction of alanine residues that ensure high helix-forming propensity ^18-19, 27^. The peptides are also highly soluble due to the high fraction of both positively and negatively charged side chains, which additionally introduce salt bridges that stabilize the helical structure ^18-19, 23^. Moreover, both glutamate and arginine are straight-chain amino acids and provide further intrinsic helix-forming propensity. Finally, the use of only non-polar sidechains in the X position (A, L, M, I, V, G) also ensures that no additional stabilizing or destabilizing polar interactions with the rest of the sequence are introduced. At the same time, these non-polar side chains span the entire range of the helix-propensity scale from the most stabilizing (alanine) to the most destabilizing (glycine) ^21-22, 26-31^.

Circular dichroism spectroscopy is one of the most powerful methods to monitor α-helical content. The molar ellipticity at 222 nm ([*Θ*]^222nm^) has traditionally been used to calculate the helical content in a system of interest ^24, 27, 30, 32-33^. Importantly, this involves the use of ellipticity values for a fully helical and fully coiled structure. Thus, to evaluate the effect of pressure on the helical content of a peptide using ellipticity measurements, one first needs to determine the pressure dependencies of ellipticities for fully helical and fully coiled structures. We have shown previously that the GEARA6 sequence remains in a coiled state at all temperatures ^18^. We thus used this peptide to record [*Θ*]_222nm_ of a fully coiled peptide as a function of pressure at three temperatures: 5°C, 20°C and 30°C (Figure 1A). The ellipticity values are very small at ambient pressure in agreement with the expectations ^18, 24, 33^ and slightly decreased with increased pressure. Based on these results, the overall equation to describe both the temperature and pressure dependence of the ellipticity of the coiled state is:

**Figure 1.**
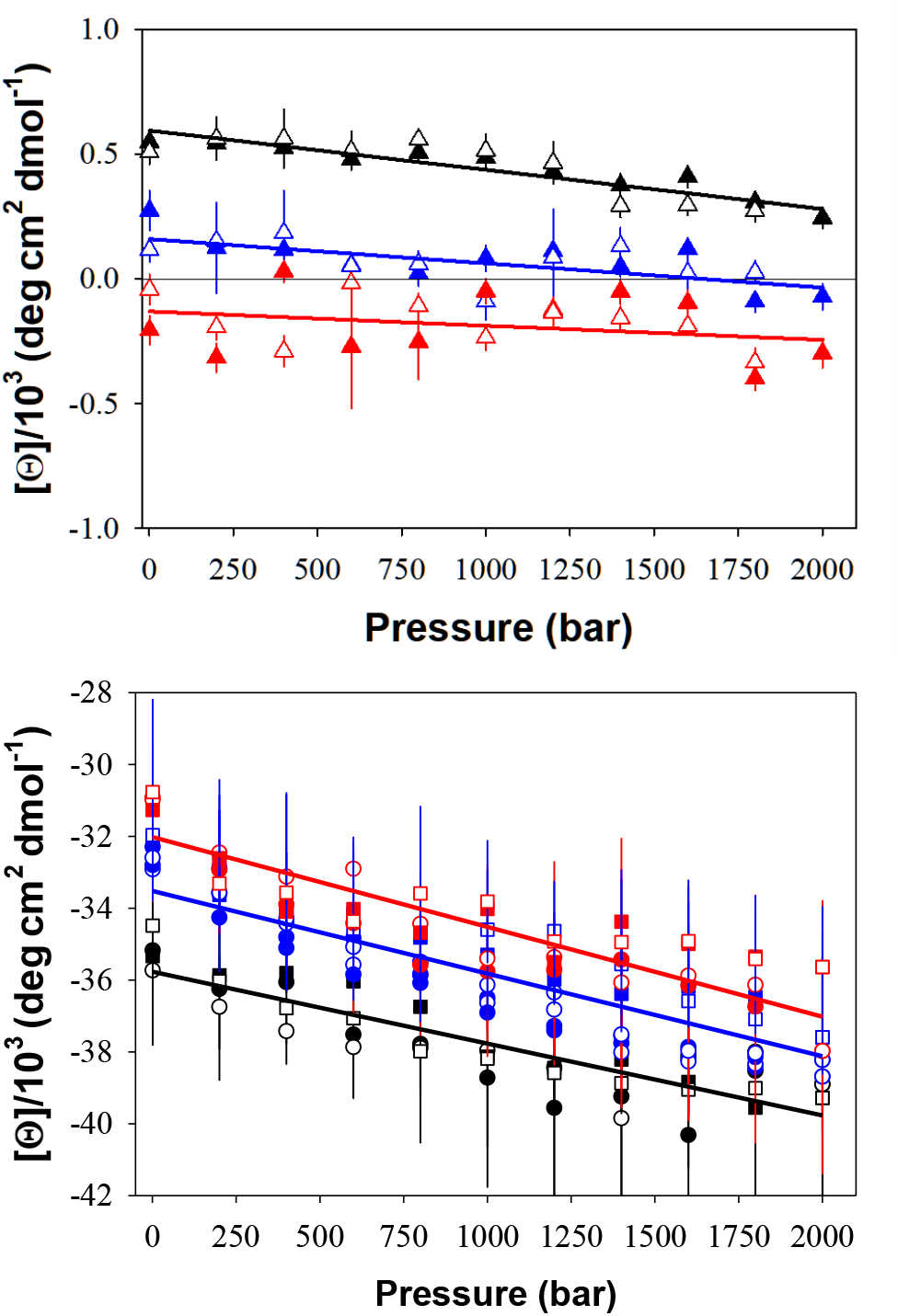
Ellipticity at 222 nm for fully coiled and fully helical structures. <u>Panel A</u>. Pressure-dependence of [*Θ*]_222nm_ at 5°C (▴), 20°C 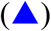 and 30°C 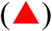 for the GEARA6 peptide. Solid lines represent the linear fit of the data according to Equation 1. <u>Panel B.</u> Pressure-dependence of [*Θ*]_222nm_ at 5°C (black), 20°C (blue), and 30°C (red) for the AEARA6 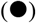 and IEARA6 (◼) peptides in 60% TFE. Solid lines represent the linear fit of the data according to Equation 2. On both panels, closed and open symbols represent the [*Θ*]_222nm_ values collected during the up and down pressure ramp, respectively, demonstrating reversibility.

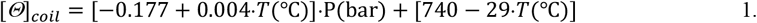

It is worth mentioning that these estimates are close to the previously established temperature dependence of the ellipticity of the coil at ambient pressure [*Θ*]_*coil*_ = 640− 45 ∗ *T*(°C) ^31^.

To experimentally access the ellipticity of a fully helical structure, we used the AEARA6 and IEARA6 peptides dissolved in a buffer containing 60% trifluoroethanol (TFE). High concentrations of TFE are known to increase the helical content of peptides with significant helical structure in water to almost 100% ^33^. The pressure-dependencies of [*Θ*]_222nm_ for AEARA6 and IEARA6 in 60% TFE buffer at a given temperature are very similar (Figure 1B) despite these two peptides having very different values of [*Θ*]_222nm_ in the absence of TFE at ambient pressure (see Figure 2 below). Thus, we can conclude that the pressure dependencies of [*Θ*]_222nm_ observed for both AEARA6 and IEARA6 in the presence of 60% TFE represent the intrinsic pressure dependence of fully helical peptides, which can be generalized using the following equation:

**Figure 2.**
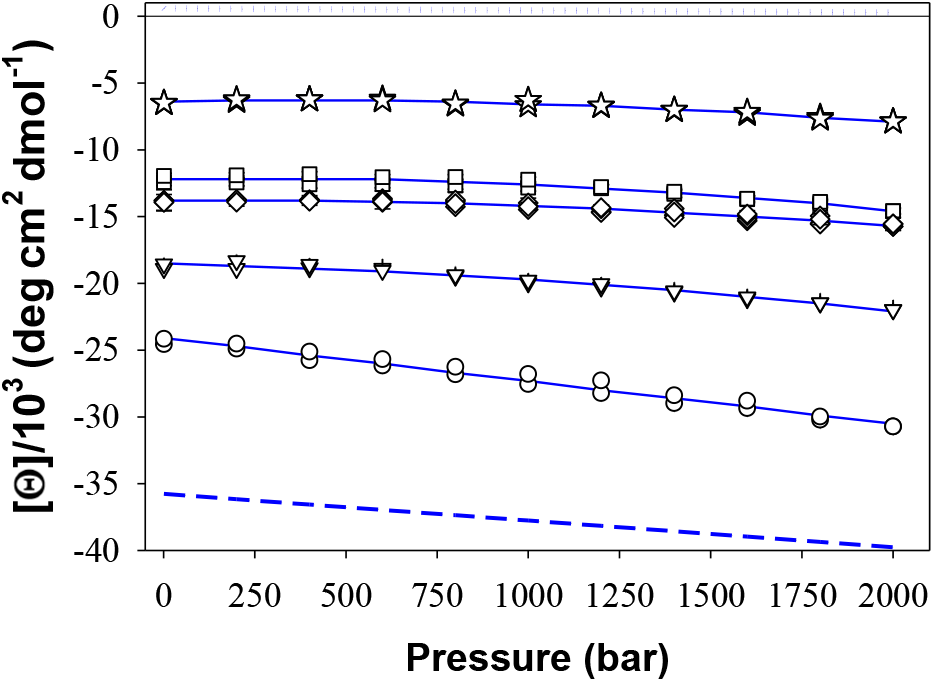
Pressure dependence of [*Θ*]_222nm_ for the AEARA6 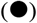, LEARA6 (▾), MEARA6 (♦), IEARA6 (◼) and VEARA6 (★) peptides at 20°C. The dotted line represents the pressure dependence of the [*Θ*]_222nm_ for the coiled state obtained from the analysis of the GEARA6 peptide, as shown in Figure 1A. The dashed line shows the pressure dependence of the [*Θ*]_222nm_ for a fully helical state obtained from the data in Figure 1B. Thin solid lines are drawn to guide the eye.

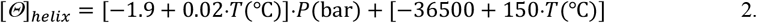

Again, these estimates are very close to the previous estimates of the temperature dependencies of helical ellipticity at ambient pressure: [*Θ*]_*helix*_ = −36800 + 230*.T*(°C) or [*Θ*]_*helix*_ = −36800 + 115*.T*(°C) ^24, 31, 33^.

Now that we have established the pressure and temperature dependencies of ellipticities for fully helical and coiled states, we can analyze the effects of pressure on the ellipticities of XEARA6 peptides in the buffer. Figure 2 compares the pressure dependence of [*Θ*]_2*2*2nm_ for five peptides, AEARA6, LEARA6, MEARA6, IEARA6, and VEARA6, at 20°C. As expected, the AEARA6 peptide has the most negative values of [*Θ*]_222nm_ at 1 bar, followed by, in order of decreasing helical propensity, LEARA6, MEARA6, IEARA6, and VEARA6 ^18^. As discussed above, GEARA6 adopts a coiled structure and is used to represent the [*Θ*]_*coil*_. The increase in pressure from ambient (1 bar) to 2000 bar causes [*Θ*]_2*2*2nm_ to become more negative. This trend observed at 20°C remains for other temperatures (see Supplementary Figure S1), albeit to a different degree depending on temperature, as is the change in [*Θ*]_222nm_ for the fully helical and coiled states (see Figure 1). Thus, to convert the readings in ellipticity [*Θ*]_222nm_ to the fraction of helical content, incorporating the [*Θ*]_*coil*_ and [*Θ*]_*helix*_ dependencies, we use the following equation:

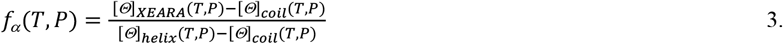

Figure 3 shows the pressure dependence of *f*_*α*_ at 20°C (for other temperatures, see Supplementary Figure S2). It is evident that helical content, in general, increases with an increase in pressure, suggesting that pressure stabilizes α-helical structures.

**Figure 3.**
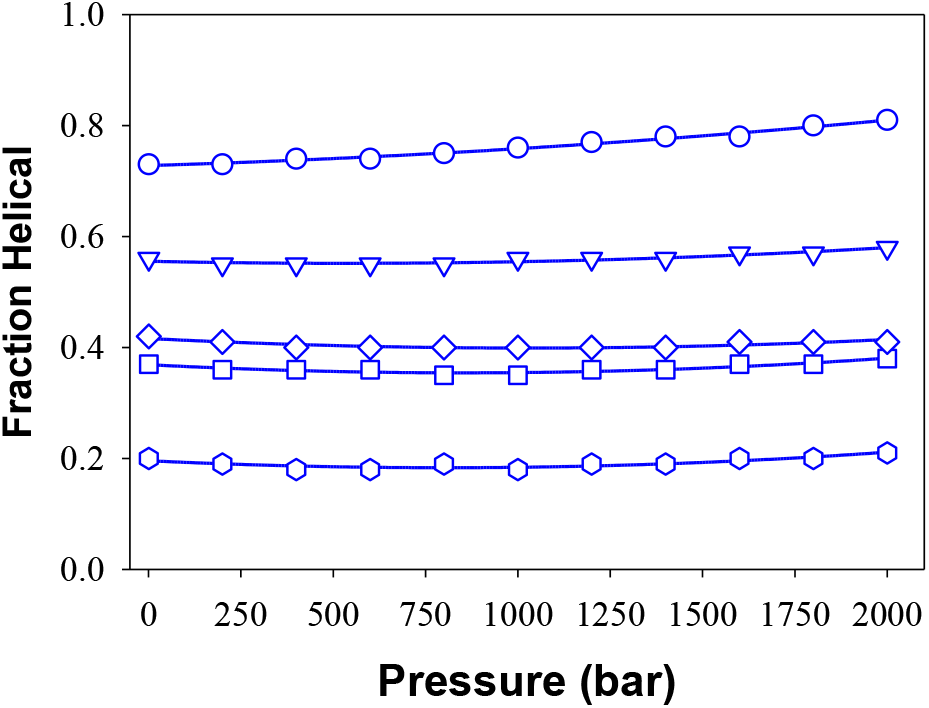
The dependence of helical fraction content, *f*_α_, on the pressure of the AEARA6 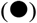, LEARA6 (▾), MEARA6 (♦), IEARA6 (◼), and VEARA6 (★) at 20°C. Values of *f*_α_ were calculated from the [*Θ*]_222nm_ at 20°C, as described by Equation 3. Thin solid lines are drawn to guide the eye.

**Figure 4.**
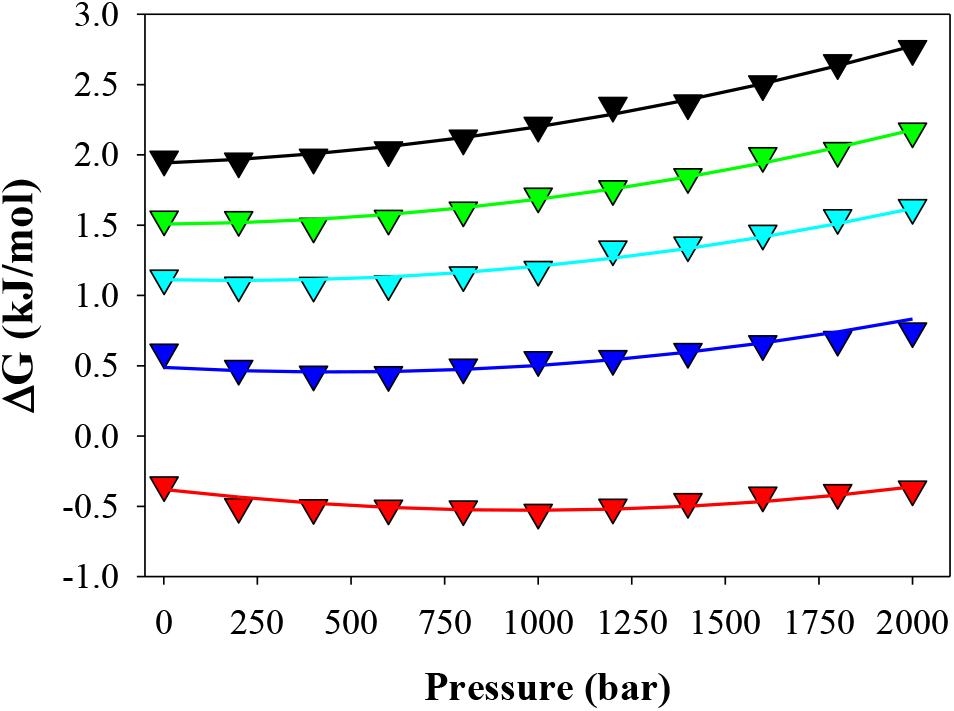
Pressure dependence of Gibbs free energy changes for the helix-coil transition of the LEARA6 peptide (see Equation 5) at 5°C (black), 10°C (green), 15°C (cyan), 20°C (blue), and 30°C (red). Solid lines represent the results of the fit to Equation 6.

According to Le Chatelier’s principle, the pressure dependence of helix stability is defined by the difference in volume between the helical and coiled states:

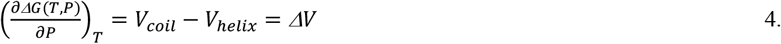

where Δ*G* is the Gibbs free energy change of the helix-coil transition, *V*_*coil*_ and *V*_*helix*_ are the volumes of the coiled and helical states, respectively, and Δ*V* is the volume change for the helix-coil transition. In a two-state approximation, the Δ*G* for the helix-coil transition can be calculated as ^18^:

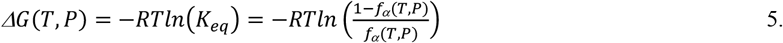

where R is the universal gas constant (8.314 J/(mol K), T is the temperature in Kelvin, and *K*_*eq*_ is the equilibrium constant. The pressure dependence of Δ*G* associated with the helix-coil transition at a given temperature can thus be described by the following equations:

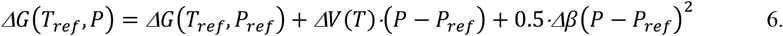

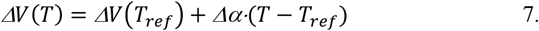

where Δα is the change in expansivity, Δβ is the change in compressibility, *ΔV* is the change in volume, and *ΔG(T*_*ref*_,*P*_*ref*_) is the Gibbs free energy change at a reference temperature and pressure (1 bar; 0.1 MPa). Data for each peptide at all temperatures were fit globally with common values for Δα, Δβ and *ΔV(T*_*ref*_). These values were found to be very similar across all peptides assessed (data for LEARA6 shown in Figure 4): Δα=-0.21±0.04 cm^3^·mol^-1^·K^-1^; Δβ=0.04±0.01 cm^3^·mol^-1^·MPa^-1^, and *ΔV(T*_*ref*_)=5.1±0.3 cm^3^·mol^-1^ (see Supplementary Figure S3 for more details on the analysis).

Imamura & Kato studied an alanine-based peptide, AK20, using FTIR spectroscopy ^16^ and found, using a two-state analysis similar to ours, positive values of ΔV: ΔV(4.5°C)=5.4±0.4 cm^3^·mol^-1^, ΔV(41.9°C)=7.2±0.6 cm^3^·mol^-1^, and ΔV(69.5°C)=8.2±0.2 cm^3^·mol^-1^. This is in good agreement with our estimates at low temperatures. However, based on the reported temperature dependence, the Δα value for the AK20 peptide appears to be positive (∼0.05 cm^3^·mol^-1^·K^-1^), which is in opposition with our estimates.

Paschek et al. also studied the effects of temperature and pressure on the thermodynamics of the AK20 peptide using molecular dynamics simulations ^14^. Their analysis determined a negative Δα=-0.9·10^−3^ cm^3^·mol^-1^·K^-1^, which agrees in sign with the Δα derived from our analysis but is somewhat larger in absolute value. The corresponding Δ*V* is very small and suggests insensitivity of the α-helical content to pressure ^14^. Furthermore, the estimates of Δβ given by Paschek et al. were between 0.016 and 0.12 cm^3^·mol^-1^·MPa^-1^, placing our value of Δβ=0.04±0.01 cm^3^·mol^-1^·MPa^-1^ well within that range.

Neumaier et al. ^15^ used a 21mer alanine-based peptide with incorporated non-natural amino acid side chains of 1-naphthyl alanine and 9-oxoxanthen-2-carboxylic acids, which allowed monitoring the triplet-triplet energy transfer between these optically active moieties as a function of pressure. Ultimately, they also found that the volume change upon helix-coil transition is accompanied by a positive ΔV=4.8 cm^3^·mol^-1^ at 5°C. Among other studies of the transitions that do not involve tertiary structure, it is worth mentioning the unfolding transition in collagen is also accompanied by positive volume changes (0.97 cm^3^·mol^-1^), shown independently by both dilatometry ^34^ and DSC ^35^.

We also applied the computational protocol we had developed previously ^10, 36^ to predict the volume changes upon unfolding of the AEARA6 peptide. The native helical state ensemble was generated via all-atom explicit solvent molecular dynamics simulations while the coiled state ensemble was generated using TraDes (see Materials and Methods for details). We found that the molecular volume occupied by the native state ensemble is smaller than the molecular volume of the unfolded state ensemble (i.e. positive volume change upon helix-coil transition). Our simulations predict a molecular volume change of 51±10 Å^3^ (35±6 cm^3^·mol^-1^), which translates to a per residue value of 1.7±0.3 Å^3^ volume increase upon helix unfolding. This estimate is identical to computational estimates done by Krobath et al. ^13^ 1.73±0.1 on a 16-residue polyalanine peptide.

To further examine the pressure dependencies using alternative methods, we turned to monitoring the global dimensions of the peptides in response to changes in pressure. First, we designed fusion constructs consisting of fluorescent proteins, sfYellow and sfCyan ^37^, connected by either AEARA6 or GEARA6 sequence linkers. These fluorescent proteins can undergo fluorescence resonance energy transfer (FRET), allowing us to monitor conformational changes that result in a change in distance between the fluorophores (see, e.g., Supplementary Figure S4). After accounting for the temperature and pressure dependence of the fluorescent protein properties themselves, the changes in the FRET signal as a function of temperature and/or pressure will be largely defined by the changes in the conformation of the linker. One convenient way of quantifying the changes in FRET signal upon perturbation is to monitor the ratiometric fluorescence signal coming from each fluorophore measured through bulk fluorescence spectroscopy (see Equation 9 and Materials and Methods section for details). Briefly, the emission spectra of the individual donor and acceptor are measured under the same conditions as the emission spectrum of the FRET construct. Then, the linear combination of emission donor and acceptor is used to fit the spectrum of the FRET construct. The ratio of the coefficients of the fit (termed D/A) is then used as a proxy for FRET efficiency: low D/A corresponds to high FRET efficiency, as the donor and acceptor get closer together, and visa-versa high D/A corresponds to a decrease in FRET as the donor and acceptor get farther apart.

Figure 5 shows D/A values for the sfYellow-GEARA6-sfCyan construct as a function of pressure at three different temperatures: 10°C, 25°C, and 40°C. The D/A profiles for different temperatures overlap, suggesting little, if any, effects of temperature or pressure on the conformational ensemble of GEARA6 linker. This is consistent with the results of CD measurement, showing that GEARA6 appears to be in a coiled state that does not show a significant change in the ellipticity at 222 nm as a function of both temperature and pressure (Figure 1). The D/A values for sfYellow-AEARA6-sfCyan construct as a function of pressure at the three different temperatures show very different trends (Figure 5). First, the D/A values at each temperature and ambient pressure are higher than those observed with the GEARA6 linker, suggesting the distance between donor and acceptor is larger; this is consistent with the AEARA6 sequence containing helical structure that will maintain a larger average end-to-end distance than the coiled GEARA6 sequence. Second, the D/A value at ambient pressure decreases with the increase of temperature from 10°C to 40°C. Again, this is consistent with the melting of helical structure in AEARA6, as has been observed by changes in the ellipticity at 222 nm (Figure 2). Third, the D/A value for the AEARA6 FRET construct at 10°C and 25°C increases with the increase in pressure, and this trend is more pronounced at 10°C. Such an increase in D/A suggests the increase in distance between the FRET pair and is consistent with the increase in the helical structure in the AEARA6 peptide upon an increase in pressure. At 40°C, the D/A for the AEARA6 FRET construct, despite maintaining an absolute value higher than that of the GEARA6 FRET construct, shows similar independence on pressure. Thermodynamic analysis of the temperature and pressure dependencies of the helix-to-coil transition in XEARA6 peptides described above shows that the changes in thermal expansivity, Δα, is negative, and thus, it is expected that the pressure stabilization of the helical structure will diminish at higher temperatures, as evidenced by the decrease in the degree of pressure-dependence with increasing temperature.

**Figure 5.**
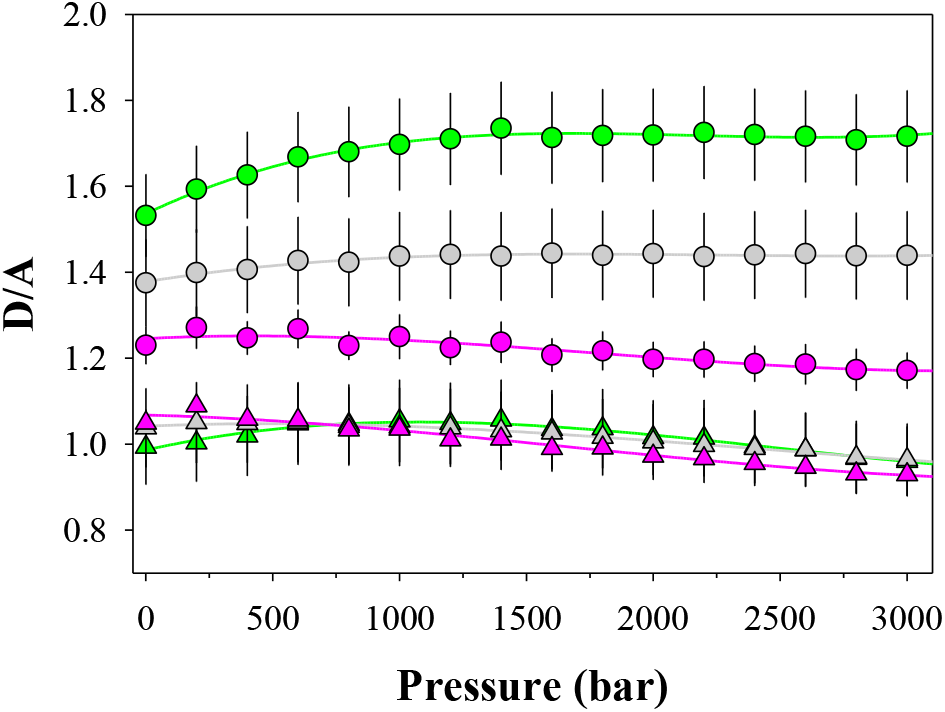
The D/A calculated for GEARA6 (▴) and AEARA6 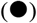 linker sequences as a function of pressure at different temperatures: 10°C (green), 25°C (gray), and 40°C (pink). D/A was calculated according to Equation 9.

Additional insight into the temperature and pressure-dependent conformational changes was obtained through orthogonal small-angle X-ray scattering (SAXS) methodology. The relatively large size of the GEARA6 and AEARA6 FRET constructs makes them suitable for SAXS experiments. The experimental SAXS profiles were analyzed using the Ensemble Optimization Method (EOM) to extract information on the conformational ensembles (described in detail in the Materials and Methods section). Using EOM, we identified a subset of 150 structural models, generated through molecular dynamics simulation, that are consistent with each experimental scattering profile (see Supplementary Figure S5). These structures were analyzed to extract the end-to-end (E2E) distance distribution of the linker regions (i.e. excluding the fluorescent protein domains). The E2E for the GEARA6 linker sequence was rather broad and was fitted to a Gaussian distribution:

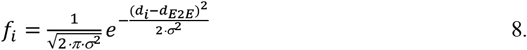

where d_E2E_ is the mean E2E distance and σ is the standard deviation of the E2E distance distribution, with d_E2E_=33±10 Å (Figure 6). This distribution is similar to the E2E distance for a coiled conformational ensemble of GEARA6 generated using TraDes (see Material and Methods for details) with d_E2E_ =32±12 Å. These support previous assessments based on the [*Θ*]_222 nm_ and D/A analysis that the GEARA6 sequence samples a coiled conformational ensemble with little to no pressure or temperature dependence. The E2E distance distribution for the AEARA6 linker sequence calculated from the EOM structures is shifted to a higher distance and follows a much narrower distribution compared to GEARA6 (Figure 6), consistent with a larger average E2E distance maintained through helical content. The AEARA6 linker within the EOM-selected models has an average d_E2E_=42±4 Å This distribution is very different from that expected for the coiled ensemble of AEARA6 generated using TraDes (d_E2E_=33±13 Å). Modeling shows that an idealized α-helix of the AEARA6 sequence (i.e. *f*_*α*_=1.0) has d_E2E_=47 Å, while molecular dynamics simulations for an ensemble with an average *f*_*α*_=0.93 has d_E2E_=45±2 Å. Calculation of the average *f*_*α*_ for AEARA6 linker from the EOM-selected models yields *f*_*α*_=0.70±0.15, so d_E2E_=42±4 Å is a reasonable estimation. It is worth noting that SAXS is an inherently low-resolution method and, while utilizing downstream analysis procedures can provide important structural information, the level of detail that can be gathered is, hence, limited in terms of precision and resolution ^38^. Therefore, although we are unable to distinguish any significant temperature or pressure-dependent factors through the analysis of SAXS data, the results of the EOM analysis are consistent with the conclusions made based on the results of CD spectroscopy and FRET data previously discussed.

**Figure 6.**
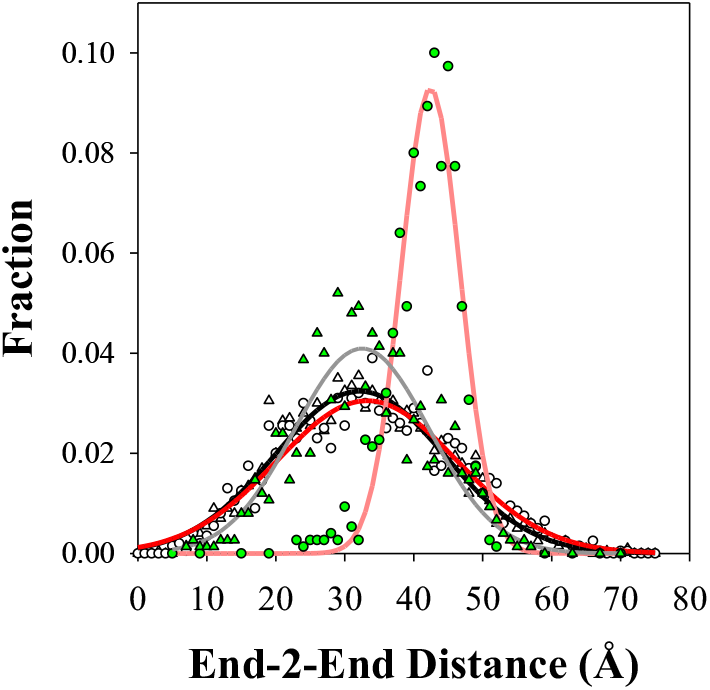
Distribution of the E2E distances for GEARA6 (green ▴) and AEARA6 (green 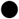) linker sequences in the EOM-selected structures (25°C). The gray and pink lines show the fit to a Gaussian distribution (Equation 7) d_E2E_(GEARA6)=33±10 Å and d_E2E_(AEARA6)=42±4 Å, respectively. Open symbols show the distribution based on the coiled state modeling using TraDes with the Gaussian fit of 32±12 Å for GEARA6 (black line) and 33±13 Å AEARA6 (red line).

## Conclusions

We find that at low temperatures, the volume changes for the helix-coil transition are positive but rapidly decrease (due to the negative temperature dependence defined by Δα), and changes sign at ∼40°C (Figure 7). At temperatures above 40°C, the helix becomes destabilized by an increase in pressure. The consequence of this is that for helical proteins with individual helices having high helical propensity, cold denaturation will lead to the formation of the highly helical denatured state, more so at high pressure. Furthermore, a molten globule state, i.e. a state without a unique tertiary structure but containing a high fraction of secondary, and in particular helical structure, will be further stabilized by high pressure and low temperatures.

**Figure 7.**
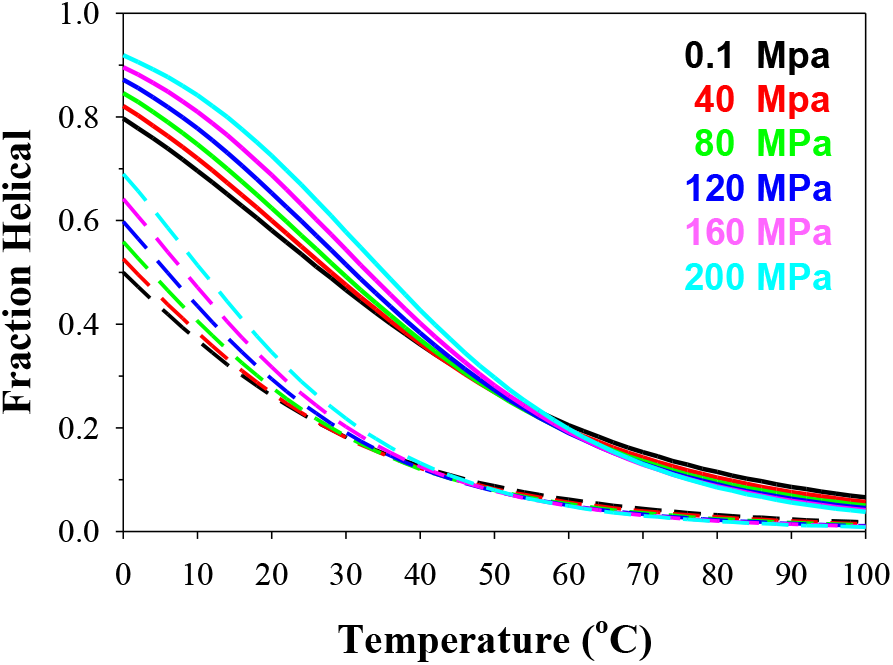
Simulated effects of the increase of pressure on the temperature-induced melting of a peptide with starting helical content of 0.8 (solid lines) and 0.5 (dashed lines). Simulations are done using Δ*V*, Δ*α* and Δ*β* parameters obtained from global analysis of all data, as described in the text.

## Materials and Methods

### Expression and purification of peptides and proteins

#### XEARA6 Peptides

The 6 peptides with six repeats of XEARA sequence (XEARA)_6_-Y, where X is A, L, M, I, V, or G were chemically synthesized, purified, and prepared for CD experiments as described before ^18-19^. The concentration of peptides was measured spectrophotometrically using the extinction coefficient *E*_1 cm;276 nm_=1,280 o.u./mol. A correction for light scattering was applied as described by Winder & Gent ^39^.

#### Fluorescent proteins

sfYellow-GEARA6-sfCyan and sfYellow-AEARA6-sfCyan were cloned into the pGia plasmid under control by the T7 promoter. Actual sequences of the proteins are given in Supplementary Table S1. The design procedure of the sfYellow and sfCyan FRET-compatible pair was described earlier ^37^. Expression and purification were done as described previously. *E. coli* BL21(DE3) cells containing the expression plasmid were grown overnight in a 5 mL starter culture of 2xYT media containing 100 μg/mL ampicillin at 37°C. 1 mL starter culture was pelleted at 1,800 x g and resuspended in fresh media, which was used to inoculate 1 L of fresh media in a baffled Fernbach flask. The culture was then incubated at 37°C while shaking at 225 rpm until OD_600_ 0.6-0.8 where expression was induced using 1 mM IPTG. The induced culture was incubated at 37°C while shaking at 225 rpm for 4-8 hr. Cells were harvested by centrifugation at 2,900 x g and 4°C for 30 min, spent media was discarded, and pellets were frozen at -20°C for >12 hr. The cell pellets were then thawed at room temperature, resuspended with ∼10 mL 20 mM Tris pH 8.0, 200 mM NaCl, 10 mM imidazole and lysed by passing twice through an ice-cold French pressure cell. Cell debris was pelleted by centrifugation of the cell lysate at 13,600 x g and 4°C for 1 hr. The supernatant was loaded to a ∼30 mL Ni^2+^-NTA column (ThermoFisher Scientific), washed with 20 mM Tris pH 8.0, 200 mM NaCl, 10 mM imidazole, and eluted using 20 mM Tris pH 8.0, 200 mM NaCl, 200 mM imidazole. The nickel column eluant was then dialyzed against 100 mM Tris pH 8.0, 5 mM EDTA using a 3.5 kDa MWCO dialysis membrane (Fisher Scientific) at 4°C overnight. The sample was loaded to a ∼30 mL DEAE Sepharose G25 Fast Flow column (Cytiva Life Sciences, Marlborough, MA) and eluted using a linear 0-500 mM NaCl gradient in 100 mM Tris pH 8.0, 5 mM EDTA, followed by size exclusion chromatography using a 2.5 × 120 cm Sephadex G-75 column (Cytiva Life Sciences) in PBS pH 7.4 with 0.02% sodium azide (will be referred to as PBSa). Fractions were checked for purity via SDS-PAGE using SurePAGE Bis-Tris, 8-16% acrylamide precast gels and Tris-MES buffer (GenScript, Piscataway, NJ), as well as MALDI-TOF. Pure fractions were pooled, concentrated using a 10 kDa MWCO centrifugal filter (Pall Corporation, Port Washington, NY), and stored at -20°C in PBSa and 20% glycerol.

#### Circular Dichroism spectroscopy

The ISS high-pressure cell has been retrofitted to fit into the Jasco-J700 circular dichroism spectrometer. The major modification was the use of MgF_2_ windows that do not undergo decrease or loss of CD signal in the far-UV at high pressures, like windows made out of quartz, and do not show pressure-induced depolarization as windows made from birefringent sapphire ^40^. Pressure is generated using a HUB440 high-pressure pump (Pressure BioSciences Inc., Medford, MA) and hydrostatic pressure is transduced to the sample using degassed deionized water. The temperature of the cell was controlled using an external water circulating water bath (refrigerated circulator model F25-ME, Julabo USA, Allentown, PA). The pressure ramp every 200 bar was done gradually over 30 sec, followed by 5 min equilibration. Ellipticity values between 221 and 223 nm every 0.2 nm at every pressure were used for calculating reported ellipticity values at 222 nm. After reaching 2000 bar, the pressure was lowered in 200 bar intervals back to 1 bar using a similar pressure change scheme, i.e., drop in pressure over 30 sec followed by equilibration for 5 min. Thus, each experiment produced similar duplicate ellipticity readings at each pressure, supporting the reversibility of pressure effects on the ellipticity signal. This was repeated at least twice for most peptides at 5, 10, 15, 20, and 30°C.

#### Fluorescence spectroscopy

Triplicate experiments were performed on a Fluoromax-4 spectrofluorometer (Horiba Jobin Yvon, Kyoto, Japan) at 25°C using a quartz microcuvette with a 10 mm path length. The sample was then excited at 430 nm (1.5 nm slit widths), and the emission spectrum scaled by wavelength-dependent lamp intensity (i.e., S/R) was detected from 440-700 nm (2 nm slit widths) with right angle geometry. Inner filter effects resulting from (1) attenuation of excitation light passing through the sample and (2) re-absorbance of emitted photons were assumed not to affect the observed ratios of donor/acceptor fluorescence (i.e., D/A, see below).

The spectra were corrected for direct acceptor excitation by subtracting the emission of sfYellow upon excitation at 430 nm from each spectrum. The spectra were then unmixed to separate sfCyan and sfYellow emissions using the following equation:

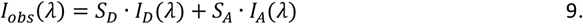

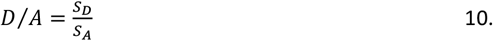

Where, at a given wavelength, *I*_*obs*_ is the observed fluorescence intensity, *I*_*D*_ and *I*_*A*_ are the reference donor (sfCyan) and acceptor (sfYellow) emission intensities, respectively, that were measured in donor- and acceptor-only samples using the same acquisition parameters (both temperature and pressure), and *S*_*D*_ and *S*_*A*_ are fitted scaling factors applied to the reference donor and acceptor fluorescence intensities such that their sum yields the observed intensity. The qualities of the fits were assessed by monitoring the residuals upon subtracting the reconstructed spectra from the observed spectra. The ratio of *S*_*D*_ to *S*_*A*_ is referred to as D/A.

#### High-pressure small-angle X-ray scattering (HP-SAXS)

HP-SAXS data were collected at the ID7A BioSAXS beamline of the Cornell High Energy Synchrotron Source (CHESS) ^41^. Briefly, 50 μL of the sample was loaded into a disposable cuvette comprised of a plastic frame encased in 0.0003” thick Kapton® HN Film (CS Hyde Company, Lake Villa, IL) and plugged using High Vacuum Grease (Dow-Corning, Midland, MI). The cuvette was then loaded into the sample cell thermoregulated by a water bath set to the desired temperature. Pressure was generated using a HUB440 high-pressure pump (Pressure BioSciences Inc.) and hydrostatic pressure was transduced to the sample using water. Isothermal pressure ramps were conducted in a buffer containing 50 mM Tris pH 7.4, 100 mM NaCl, and 5% glycerol. Samples for HP-SAXS were concentrated to the highest allowable concentration, avoiding aggregation. Immediately prior to SAXS data collection, the samples were centrifuged for 10 minutes at room temperature to remove any insoluble particles. 5 exposures of 1 second each were recorded at each pressure, ranging from 50-3000 bar. Buffer subtraction and Guinier analysis were conducted using the BioXTAS RAW software (v2.2.1) ^42^ to assess data quality. The lowest q values that significantly deviated from linearity in the Guinier region due to higher order structures and/or artifacts near the beam stop were truncated from the experimental profile.

The ensemble optimization method (EOM) program suite (v3.0) ^38, 43^ was used to fit the experimental SAXS profiles to theoretical scattering patterns generated by an ensemble of conformers. Structural models of each fusion construct were generated using AlphaFold2 (v2.12) ^44^. All-atom structure-based potentials were generated from these models using the SMOG web server (v.2.0.3) with default parameter sets ^45-46^. The contacts were identified from PDB coordinates through the use of the Shadow Contact Map algorithm ^47^ with a cutoff distance of 6 Å, shadowing radius of 1 Å, and residue sequence separations of 3. Atom pairs that are not identified as contacts were assigned an excluded volume interaction. The bond lengths and angles and improper and planar dihedral angles of the protein were maintained by harmonic potentials. The potentials were assigned such that the native configuration of each bond and angle was considered the minimum. The final form of the potential energy function for the all-atom structure-based model is:

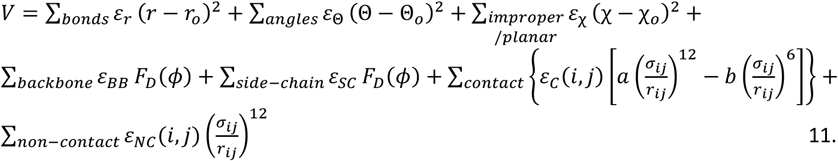

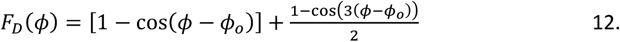

with all parameters having the default values as reported in ^45^. The potentials were further modified to increase the strength of Lennard-Jones interactions within the sfYellow and sfCyan domains by a factor of 1.5 to ensure that the structure of the fluorescent protein domains remained unchanged at all temperatures during the simulations. Furthermore, any Lennard-Jones interactions in the GEARA6 linker sequence were removed to ensure that no specific constraints in this region were enforced during the simulation. GROMACS (v.4.6.7) ^48^ was used as the computation engine to run the simulations. Simulations using Langevin dynamics (time step, τ = 0.0005 ps) were run for 5·10^8^ steps at both low (110K) and high (130K) temperatures to ensure sufficient coverage of the conformational space. 5,000 frames from each of the trajectories were extracted at random (10,000 total) to yield the final pool of structural models. The models were assessed manually and were determined to adequately span the allowable conformational space of each construct.

The theoretical scattering intensities and size statistics of each model were then computed by FFMAKER for input into the Genetic Algorithm Judging Optimization of Ensembles (GAJOE), which employs a genetic algorithm approach to select an ensemble of models that best describes the experimental scattering profile ^38, 43^. GAJOE was run 3 independent times for 200 cycles of 5,000 generations each, with the number of ensembles per generation set to 50. The number of theoretical curves in each ensemble was fixed at 50 while also disallowing curve repetition; these parameters were selected based on assessments made by the software developers that analysis, where the ensemble size is optimized, may yield artificially bimodal distributions of flexible systems ^49^. R_flex_ and R_σ_, metrics that describe the flexibility of the selected ensemble both independently and relative to the flexibility of the starting pool, respectively, were taken together to assess the fit quality as described by the developers ^38^. Theoretical scattering curves were calculated for the ensemble of 150 structures using CRYSOL ^50^ and averaged for comparison to the experimental profiles. The *χ* ^2^ value reporting on the goodness of fit of the averaged theoretical scattering profiles to the experimental profile was calculated from the equations outlined by the developers ^50^. The secondary structural content of the linkers in the ensembles for each temperature/pressure condition was assessed using DSSP ^51^.

#### Other computational methods

The starting structure of the AEARA6 sequence in PDB format adopting the ideal α-helix was generated using SwissPDB ^52^. This PDB file was used for all-atom explicit solvent simulation using the charmm36 forcefield ^53^ and tip3p water model. Molecular dynamics simulations were carried out in GROMACS 2022.2 ^48^. The native crystal structure was solvated in a dodecahedron box, with dimensions such that all protein atoms are at least 10 Å deep in the box, and neutralized with 0.1M excess NaCl, followed by energy minimization for 1,000 steps. All simulations underwent 200 ps of constant volume equilibration, 200 ps of constant pressure equilibration and 200 ns of production simulation at a defined temperature and pressure and 1 bar. We used the Parrinello–Rahman pressure control ^54^ with a 2 ps relaxation time, and a compressibility of 4.6×10^−5^ atm^-1^, and v-rescale temperature coupling. LINCS ^55^ and SETTLE ^56^ algorithms were used to constrain high-frequency bond vibrations that allowed the use of a 2 ps integration step. The electrostatic interactions were modeled by the smooth particle mesh Ewald method ^57^, using a 75×75×75 grid, with fourth-order charge interpolation. A native ensemble of 200 structures was extracted from the production trajectory (1 structure per ns). The persistence of helical structure throughout the simulations was validated by monitoring the backbone dihedral angles using the g_rama utility from GROMACS ^48^.

Sampling of the unfolded state ensembles was carried out using TraDes ^58^. TraDes ensembles were generated as described before ^10^ and consisted of 2000 structures. Before volume calculations, the structures generated by TraDes were energy minimized in implicit solvent, using the Generalized Born surface area (GBSA) model with protein and solvent dielectric constants of 80 in GROMACS 4.6.7. This step also explicitly incorporates all hydrogen atoms.

The ProteinVolume ^36^ and MSMS ^59^ software packages were used to calculate the volume and molecular surfaces for each structure in the native or unfolded ensembles, and ensemble-averaged values were used to calculate the volume changes upon helix-coil transition as described by us earlier ^10^.

## Acknowledgments

We thank Drs. Qingqiu Huang and Richard Gillilan for help with SAXS experiments, and Dr. Tejaswi Koduru for help with using EOM.

## Supplementary Information: Figures S1-S5 and Table S1

**Figure S1.**
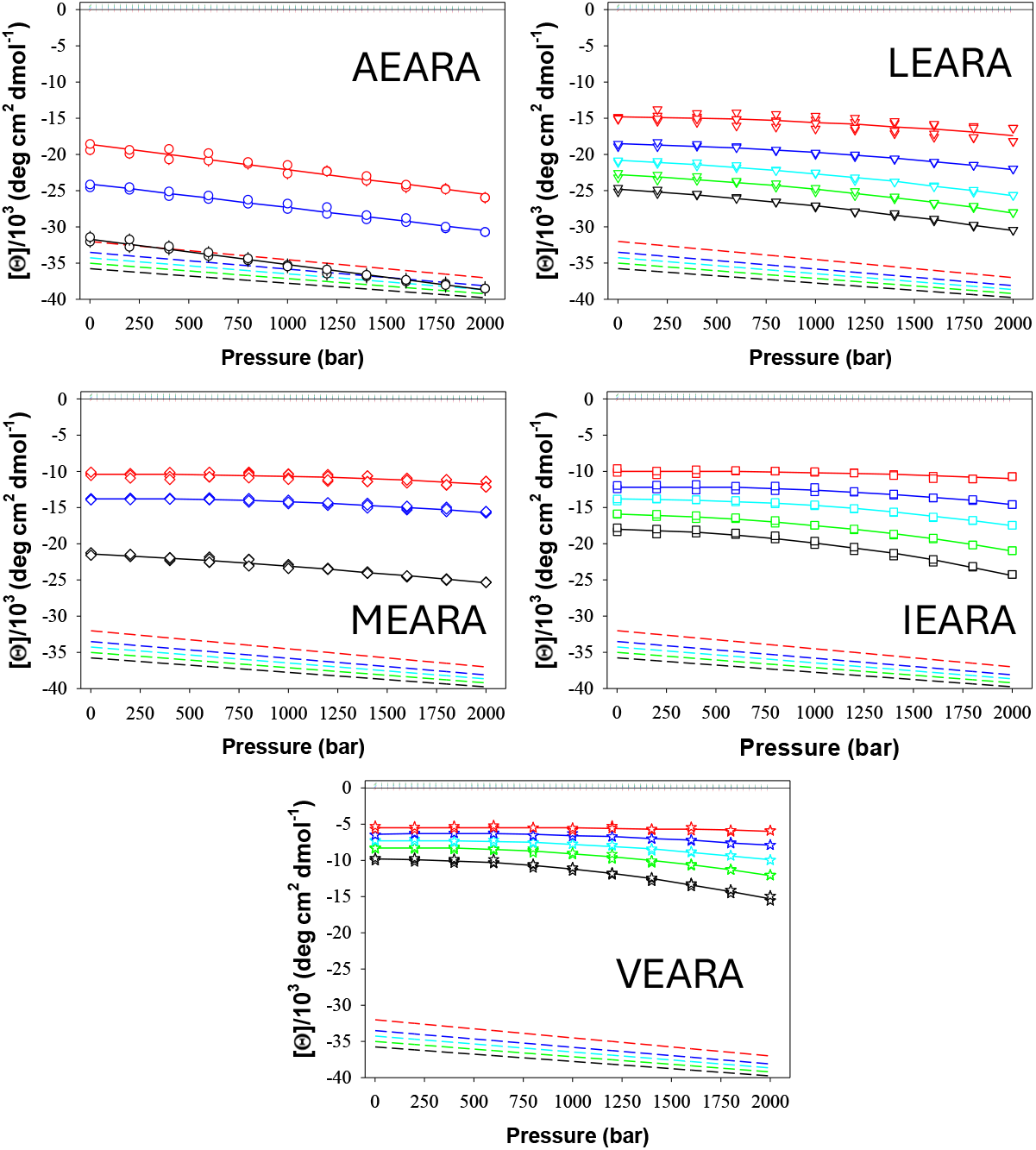
Pressure dependence of [*Θ*]_222nm_ for AEARA6 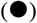, LEARA6 (▾), MEARA6 (♦), IEARA6 (◼), and VEARA6 (★) at different temperatures: 5°C (black), 10°C (green), 15°C (cyan), 20°C (blue), and 30°C (red). The dotted lines represent the pressure dependence of [*Θ*]_222nm_ for the coiled state obtained from the analysis of the GEARA6 peptide, as shown in Figure 1A. The dashed lines show the pressure dependence of [*Θ*]_222nm_ for a fully helical state obtained from the data in Figure 1B. Thin solid lines are drawn to guide the eye.

**Figure S2.**
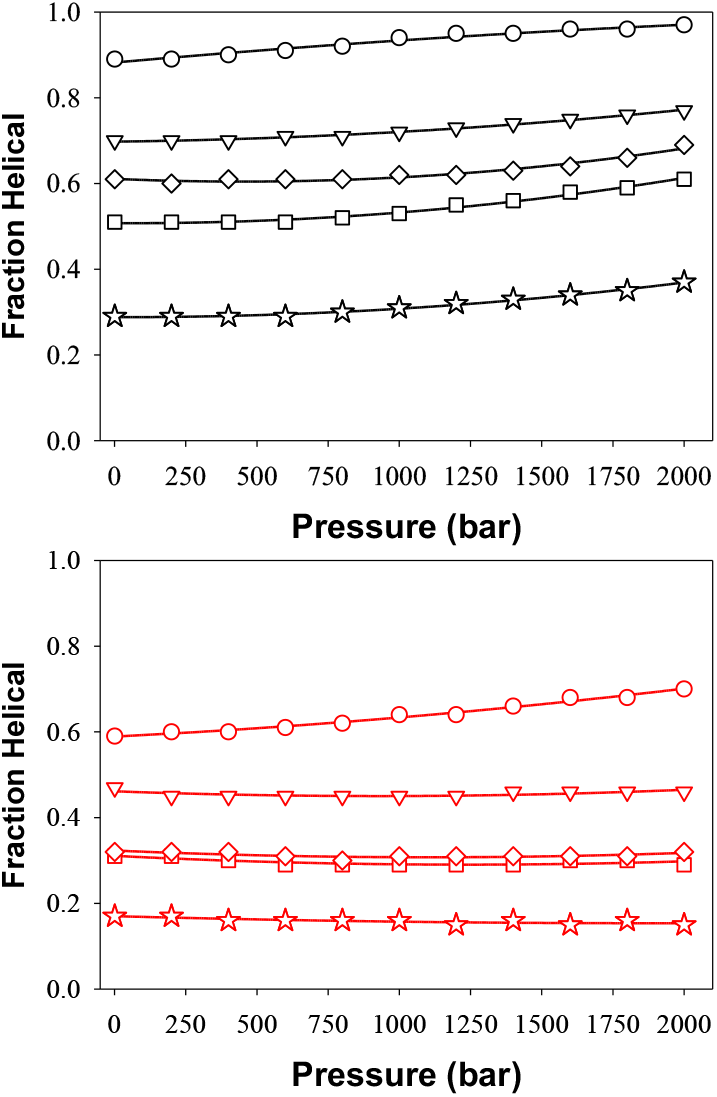
Pressure dependence of the fraction of helical content, *f*_α_, of AEARA6 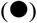, LEARA6 (▾), MEARA6 (♦), IEARA6 (◼), and VEARA6 (★) at 5°C (black), and 30°C (red). Values of *f*_α_ were calculated as described by Equation 3. Thin solid lines are drawn to guide the eye.

**Figure S3.**
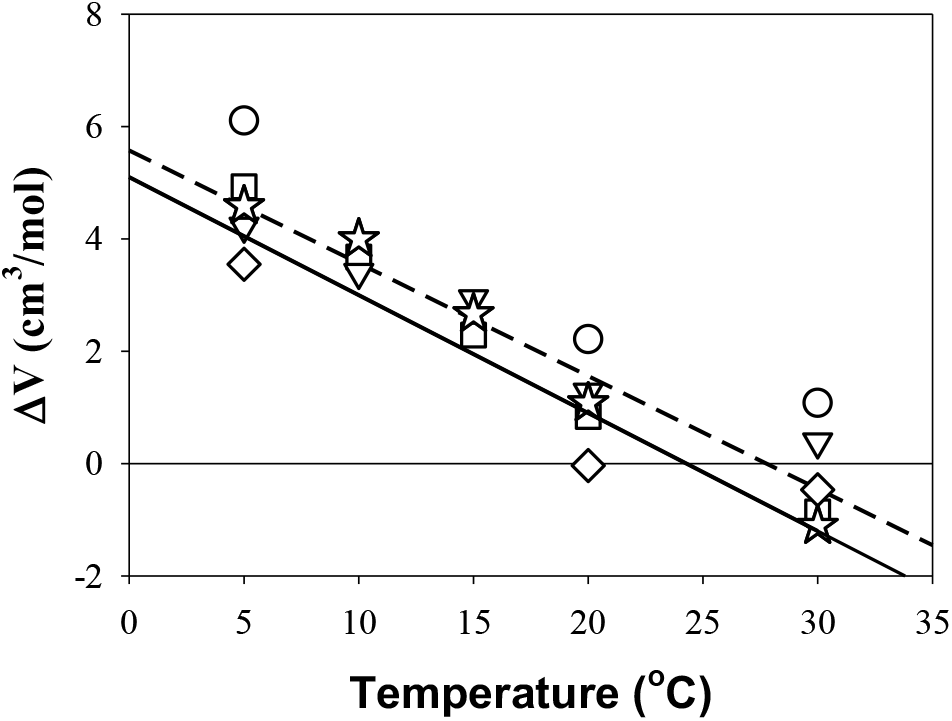
Temperature dependence of the volume change (ΔV) for the helix-coil transition. Symbols are the results of fits of ΔG versus pressure at a given temperature, while the dashed line is the fit of the data points to a straight line. The solid line is the result of global fit at all temperatures with Δ*V(T*_*ref*_)=5.1±0.3 cm^3^·mol^-1^ and Δ*α*=-0.21±0.04 cm^3^·mol^-1^·K^-1^. Symbols are for different peptides: AEARA6 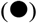, LEARA6 (▾), MEARA6 (♦), IEARA6 (◼), and VEARA6 (★).

**Figure S4.**
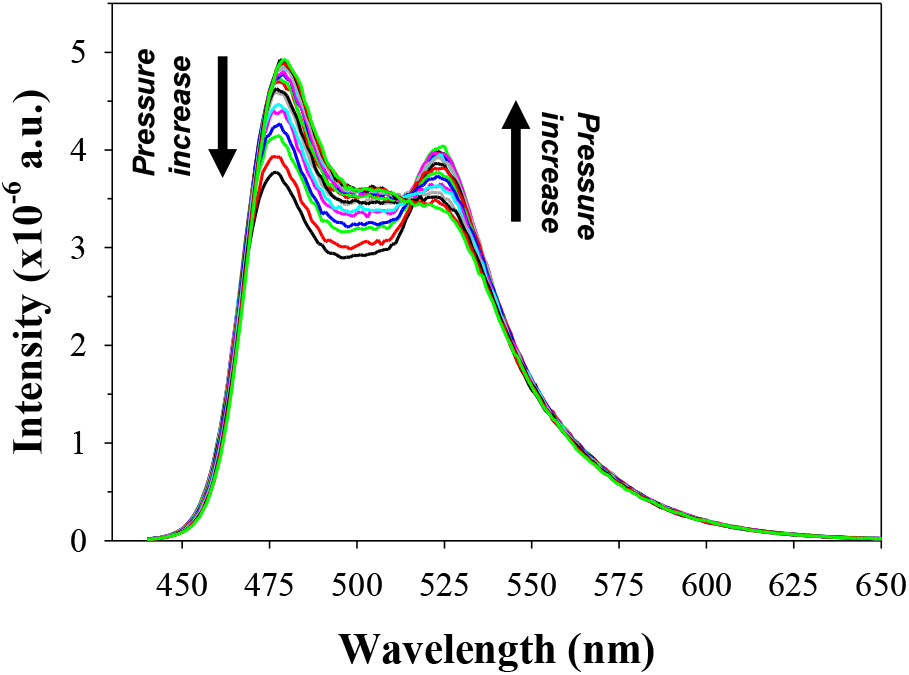
The changes in the emission spectrum of sfYellow-AEARA6-sfCyan construct at 25°C as a function of the increase in pressure from ambient to 3200 bar in 200 bar increments.

**Figure S5.**
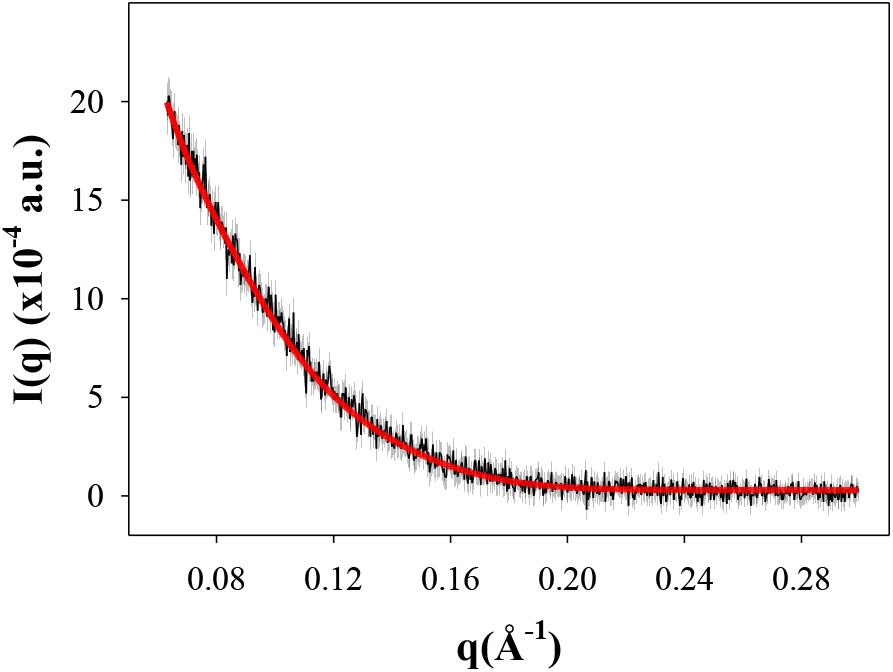
Example of fit (red) of the average theoretical scattering profile calculated by CRYSOL from the 150 EOM-selected structural models of sfYellow-AEARA6-sfCyan at 25°C at 5 MPa (experimental scattering profile shown in black).

**Table S1.**
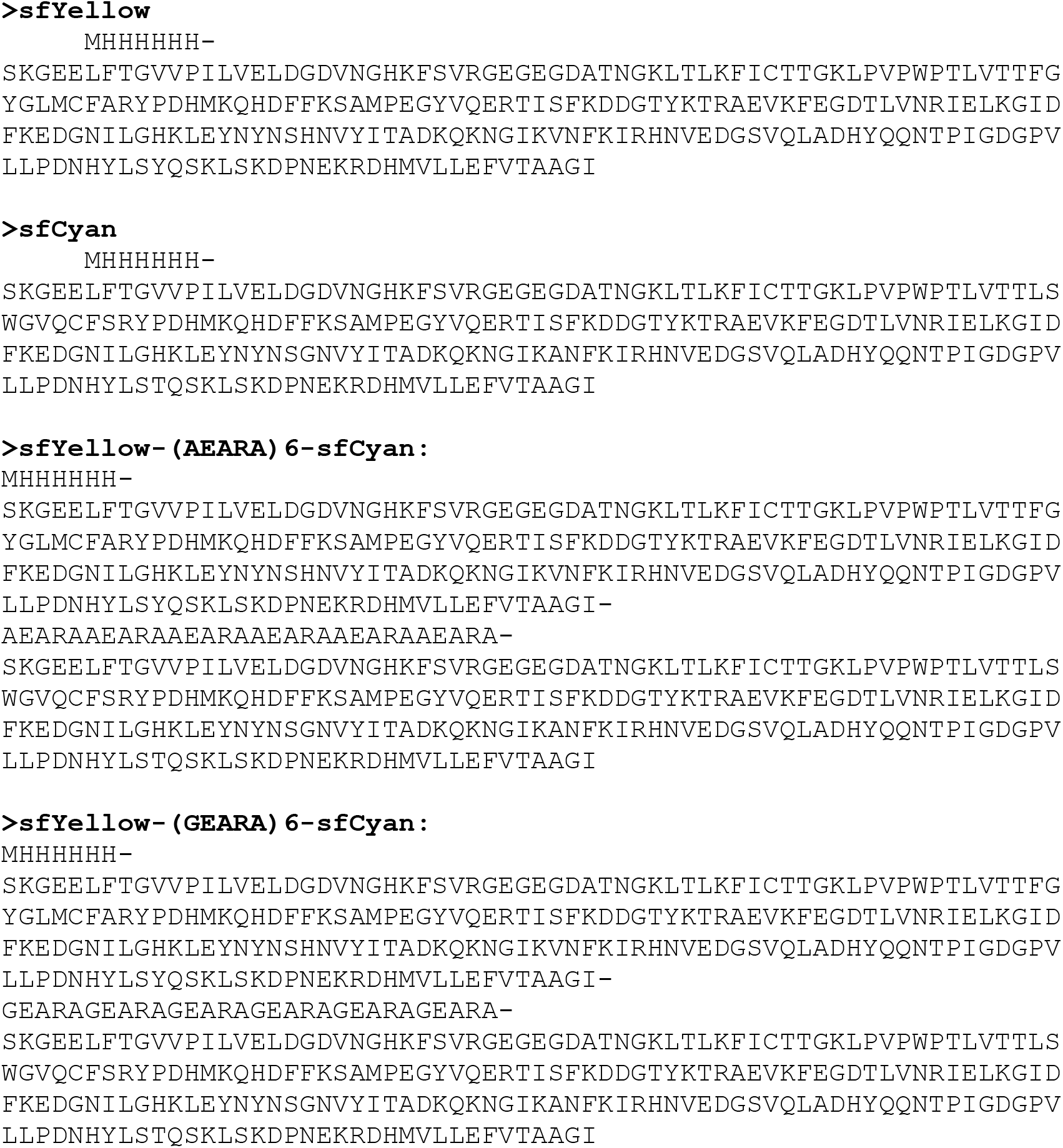
Sequences of the Proteins.

